# ProTrek: Navigating the Protein Universe through Tri-Modal Contrastive Learning

**DOI:** 10.1101/2024.05.30.596740

**Authors:** Jin Su, Xibin Zhou, Xuting Zhang, Fajie Yuan

## Abstract

ProTrek redefines protein exploration by seamlessly fusing sequence, structure, and natural language function (SSF) into an advanced tri-modal language model. Through contrastive learning, ProTrek bridges the gap between protein data and human understanding, enabling lightning-fast searches across nine SSF pairwise modality combinations. Trained on vastly larger datasets, ProTrek demonstrates quantum leaps in performance: (1) Elevating protein sequence-function interconversion by 30-60 fold; (2) Surpassing current alignment tools (i.e., Foldseek and MMseqs2) in both speed (100-fold acceleration) and accuracy, identifying functionally similar proteins with diverse structures; and (3) Outperforming ESM-2 in 9 of 11 downstream prediction tasks, setting new benchmarks in protein intelligence. These results suggest that ProTrek will become a core tool for protein searching, understanding, and analysis.

---

Proteins, the fundamental molecular machines of life, orchestrate a vast array of biological processes. Deciphering their intricate structures, diverse functions, and complex interactions lies at the heart of modern biochemistry, molecular biology, and pharmacological innovation. Biologists are driven to unravel the tripartite complexity of a protein’s sequence, structure, and function (SSF)—a molecular enigma that holds the key to unlocking the fundamental principles of life.

To address this challenge, the protein analysis landscape has evolved significantly, employing various computational approaches. Alignment-based tools like BLAST [3], MUSCLE [12] MMseqs2 [26], TM-align [37] and Foldseek [32] have made substantial contributions. However, these tools often focus on local alignment for efficiency, potentially overlooking global insights, and typically operate within a single modality—either sequence or structure—limiting their capacity to align multiple modalities concurrently. This limitation is further exacerbated by the fact that approximately 30% of proteins in UniProt [28] remain unannotated, primarily due to their sequences being phylogenetically distant from known functional homologs [4, 14]. The advent of neural network-based functional annotation tools [4, 38, 25] has ushered in a new era of protein characterization, enabling the identification of corresponding annotations for given proteins. Nevertheless, these annotation tools, primarily reliant on predefined labels, lack the sophistication to comprehend natural language, thereby impeding their ability to generate nuanced textual descriptions of protein functions or identify proteins from extensive textual narratives.

Recent breakthroughs in large language models, exemplified by ChatGPT [1] and LlaMA [30, 31], have demonstrated unprecedented capabilities across various natural language processing tasks. Concurrently, the field of computational biology has witnessed the emergence of protein language models (PLM) as a burgeoning area of exploration [19, 13, 2, 21, 36, 22]. Building upon these advancements, we posit the feasibility of establishing a foundational protein model capable of comprehensively representing protein SSF through sophisticated language modeling techniques.

Here, we introduce ProTrek, an innovative tri-modal PLM that jointly models protein SSF modalities. ProTrek leverages contrastive learning [24] through three core alignment training strategies (Fig. 1a): (1) bidirectional supervision between protein structure and sequence, (2) reciprocal supervision between protein functions and structures, and (3) mutual supervision between protein functions and sequences. This trimodal alignment training paradigm enables ProTrek to forge tight associations among SSF by converging genuine sample pairs (sequence-structure, structure-function, and sequence-function) while simultaneously diverging negative samples within the latent space.

**Fig 1.**
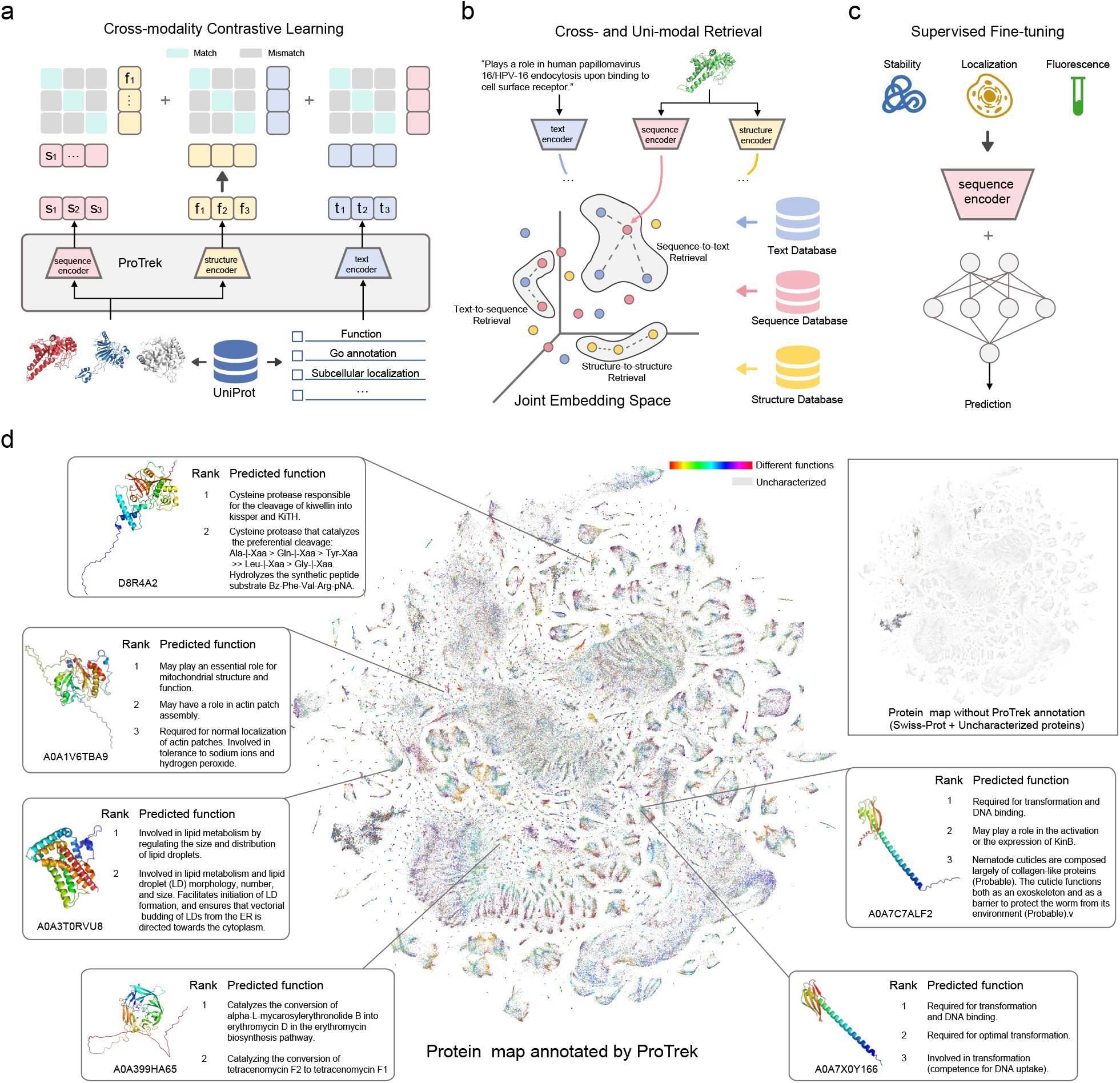
Illustration of ProTrek. **a**, ProTrek architecture and tri-modal contrast learning. **b**, Cross-modal and uni-modal retrieval. ProTrek supports nine searching tasks. **c**, After the tri-modal contrast learning, the protein sequence encoder encodes almost universal representation of proteins, which can be fine-tuned to predict diverse downstream tasks, such as protein fitness and stability prediction. **d**, Using ProTrek’s natural language capabilities to decode the protein universe. Each cluster represents proteins with close embedding distances. Over 99% of protein entries in UniProt remain unreviewed, as shown in the top right.

The architecture of ProTrek incorporates a pre-trained ESM encoder [19] for amino acid (AA) sequence modeling and a pre-trained BERT encoder [15] for nuanced natural language function representation. Protein structures are also transformed into discrete-token sequences utilizing Foldseek [32], facilitating their encoding via a BERT-style sequential network. That is, ProTrek ensures that each modality of SSF is modeled by a dedicated language model, creating a harmonious tripartite representation.

ProTrek was trained on a dataset comprising 40 million examples (see Supplementary Table 1 and 2), the largest collection of protein-text pairs at preprint release, including 14 million precise protein-text pairs curated from Swiss-Prot [5] database and 25 million relatively noisy pairs meticulously filtered from 300 million UniRef50 [7] samples (also see Supplementary Fig. 1). The ProTrek model’s training regimen incorporates 8 loss functions—6 dedicated to inter-modal alignment and 2 masked language modeling losses to preserve recognition fidelity at both the amino acid and structural (a.k.a. 3Di) token levels (see Methods).

Through an exquisite fusion of protein SSF modalities into a unified framework, ProTrek offers three pivotal capabilities that redefine our approach to decoding the protein universe: (1) As a zero-shot retrieval model, ProTrek enables precise exploration of intricate SSF interrelationships through *all* nine distinct search tasks (Fig. 1b), namely, sequence-to-structure, sequence-to-function, sequence-to-sequence, structure-to-structure, structure-to-sequence, structure-to-function, function-to-function, function-to-sequence, and function-to-structure retrieval. Notably, it bridges the chasm between protein data and human comprehension, translating complex molecular landscapes into an intuitive, language-driven experience (Fig. 1d). This paradigm shift empowers researchers with unprecedented access to nuanced natural language descriptions of protein properties and functions. (2) Harnessing the power of ‘global’ representation learning, ProTrek effectively overcomes the ‘local’ constraints inherent in current sequence comparison tools. This enables the identification of proteins with convergent functions despite divergent structures and sequences (Fig. 2b)—a phenomenon potentially more ubiquitous in nature [18, 29]. (3) Through SSF cross-modal contrastive learning, ProTrek injects structural and functional information into amino acid sequences, thereby catalyzing effective transfer learning and enabling fine-tuning across a myriad of downstream tasks, rivaling and complementing leading protein language models(Fig. 1c,2e). This versatility extends its applicability across diverse domains of protein science.

**Fig 2.**
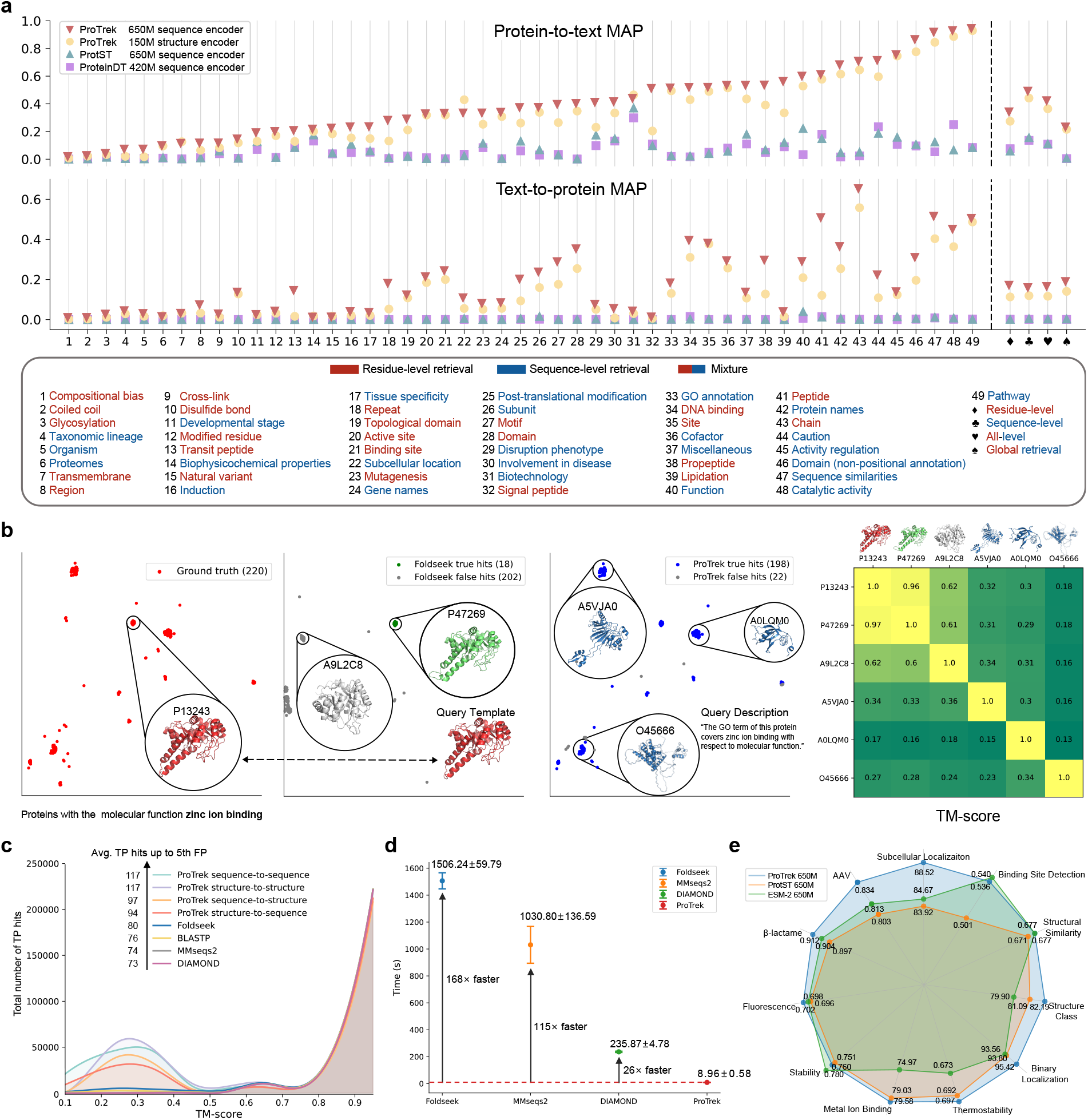
ProTrek performance on protein search and representation tasks. **a**, Top chart: Search protein functional descriptions using sequences/structures. Bottom chart: Search protein sequences/structures using textual descriptions. x-axis: Specific protein function categories left of the dashed line; aggregated categories (residue-, protein-, all-level) right of it. “Global retrieval” indicates a search across the entire database, not within individual categories. The y-axis is MAP (mean average precision), a commonly used ranking metric for searching tasks. **b**, ProTrek employs “zinc ion binding” as the query term, while Foldseek utilizes P13243 as a query template, which is the protein with the most hits. In the testing set, 220 proteins share similar functional annotations with P13243. Foldseek identified 18 true hits, whereas ProTrek discovered 198 true hits. The TM-score results in the right subfigure reveal that proteins with similar functions can exhibit diverse structures. Conversely, proteins with similar structures (e.g., A9L2CB) may encode different functions. **c**, Searching proteins with similar functions using protein sequence/structure as input. TP (true-positive): Matches sharing ≥1 GO term. FP: Matches sharing no GO terms. **d**, Comparing alignment speed (CPU time) for 100 query proteins on UniRef50 with 50 million candidate proteins, utilizing 24 CPU cores. **e**, Evaluating the protein representation ability of the ProTrek AA sequence encoder.

ProTrek is available in two variants: a large-scale version featuring a 650M AA sequence encoder, 150M structure encoder, and 130M function encoder; and a more compact version with a 35M sequence encoder, 35M structure encoder, and 130M function encoder. These configurations are designed to accommodate varying computational resources and research objectives. During inference, ProTrek employs a max-inner product search (MIPS) algorithm [11] for efficient retrieval. This enables ProTrek to complete searches within seconds, even when querying databases containing billions of entries. Such rapid search capability significantly enhances the efficiency of protein research, providing a powerful tool for exploring the protein universe.

Here, we firstly evaluated ProTrek’s performance on four emerging retrieval tasks, designed to bridge protein sequences/structures with their natural language functional descriptions. These tasks comprise bidirectional pairwise searches: sequence-to-text, structure-to-text, text-to-sequence, and text-to-structure (Fig. 2a). ProTrek’s performance was benchmarked against two state-of-the-art methods, ProteinDT (also known as ProteinCLAP)[20] and ProtST [35], using the human-reviewed Swiss-Prot [5] dataset. This evaluation dataset comprises 4,000 proteins and their associated functional descriptions. For a fair evaluation, proteins sampled in the test set have less than 50% sequence identity with those in the training set. Additionally, we incorporated 100,000 randomly sampled proteins from UniProt as unknown negative examples to further assess ProTrek’s generalization ability (see Methods).

Notably, ProTrek demonstrates superior performance across most functional categories, regardless of whether the AA sequence encoder or 3Di structure encoder is employed. It greatly outperforms both ProteinDT and ProtST, achieving remarkable improvements of over 30-fold and 60-fold in global retrieval tasks. Specifically, ProTrek attains scores of 0.233 in sequence-to-text and 0.190 in text-to-sequence search tasks, compared to ProtST’s 0.007 and 0.001, and ProteinDT’s 0.004 and 0.003, respectively (Fig. 2a). This exceptional performance can be attributed to ProTrek’s extensive training dataset, which surpasses those of ProteinDT and ProtST by two orders of magnitude. Beyond basic search capabilities, this substantial advantage positions ProTrek as a highly promising biological tool with potential applications in text-based protein design and protein question-answering systems [9, 20, 34, 33]. Within ProTrek’s architecture, the AA sequence encoder generally exhibits superior performance compared to the structure encoder, a discrepancy potentially linked to differences in encoder model size and pre-training status. Fig. 1d illustrates an example of how ProTrek embeddings and its natural language understanding capabilities can be utilized to decipher the vast protein universe.

Secondly, we assessed ProTrek’s capability in searching functionally similar proteins, benchmarking it against several established alignment-based tools, including MMseqs2 [26], DIAMOND [6], BLASTP [3], and the cutting-edge structure aligner, Foldseek [32] (Fig. 2b,c). While these classical methods rely on local alignment through sequence or structure similarity, ProTrek employs a distinctive ‘global’ alignment approach via cross-modal contrastive learning. To illustrate the difference, we evaluated the functions of retrieved proteins across four search tasks: two cross-modal searches bridging sequences and structures (sequence-to-structure and structure-to-sequence), and two uni-modal searches within sequences (sequence-to-sequence) and within structures (structure-to-structure). We conducted an exhaustive all-versus-all search on the Swiss-Prot test dataset, comprising 4,000 proteins, and compared the performance of each method in identifying proteins sharing the same GO annotation [23] up to the 5th FP (false-positive) (see Methods).

ProTrek outperforms all these classical sequence alignment tools in terms of the total/average number of correct hits (Fig. 2c). While all methods can readily identify functionally similar proteins when they share higher TM-scores with query proteins, ProTrek’s advantage becomes more pronounced when identifying functionally similar proteins with lower TM-scores. This is because proteins with similar functions can display distinct sequences or structures (Fig. 2b), as also exemplified by numerous isozymes [17] and functionally convergent proteins found in nature. Regarding ProTrek’s four search modalities, sequence-to-sequence search exhibits the highest efficacy, followed by structure-to-structure and sequence-to-structure searches. Each search task, however, offers distinct advantages. For example, while the structure-to-sequence search shows slightly lower effectiveness, it has potential applications in filtering or ranking protein sequences generated by AI-based protein design models (such as ProteinMPNN [10]). This versatility underscores ProTrek’s comprehensive approach to protein analysis and its potential to enhance various aspects of protein research and design.

ProTrek’s inference speed, bolstered by the max-inner product search (MIPS) algorithm, represents another significant advantage. This algorithmic prowess enables ProTrek to execute lightning-fast searches and sorting operations on billion-scale databases within seconds, outpacing Foldseek and MMseqs2 by over two orders of magnitude (Fig. 2d). Correspondingly, it demonstrates a speed enhancement of approximately 400,000-fold compared to TM-align [37] and Dali [16] (as reported in [32]). This exceptional search velocity empowers researchers to swiftly navigate and analyze the vast protein universe with unprecedented efficiency. Furthermore, ProTrek’s unique cross-modal search capabilities set it apart from conventional alignment tools, underscoring its pivotal role as a transformative addition to the bioinformatics toolkit.

Beyond its exceptional search capabilities, ProTrek’s amino acid and 3Di sequence encoders serve as universal representation models, owing to two unsupervised loss functions: contrastive learning and masked language modeling. We rigorously evaluated these models across a diverse spectrum of over 10 downstream protein tasks, encompassing both protein-level and residue-level analyses, as well as regression and classification challenges. Through downstream supervised fine-tuning, ProTrek demonstrated superior performance, surpassing the established ESM-2 model and the state-of-the-art ProtST model in 9 out of 11 tasks (Fig. 2e). Notably, in several tasks, ProTrek exhibited a remarkable performance boost, exceeding ESM-2 or ProtST by relative margins of 5% to 7%. These results underscore ProTrek’s robust transfer learning capabilities and its potential as a powerful representation model for a wide array of protein-related applications.

We have made the ProTrek model weights publicly available (https://huggingface.co/westlake-repl/ProTrek_650M_UniRef50) and developed a web server^1^ (https://huggingface.co/spaces/westlake-repl/Demo_ProTrek_650M_UniRef50) for multi-database searches across Swiss-Prot, AlphaFoldDB, UniRef50, and PDB. We’ve also released protein sequence embeddings for all proteins within these databases. To enhance accessibility, we’ve created a Google Colab implementation of ProTrek (https://colab.research.google.com/github/westlake-repl/SaprotHub/blob/main/colab/ColabProTrek.ipynb) for biologist-friendly fine-tuning on new data, inspired by the OPMC concept in [27].

ProTrek’s high accuracy and efficiency in cross-modal and uni-modal searches make it a valuable tool for protein database analysis, empowering biologists to navigate the vast protein universe more effectively. Its sophisticated natural language understanding capabilities transcend traditional keyword-matching limitations, enabling nuanced and context-aware searches (see Supplementary Fig.4). As a robust foundational model, ProTrek facilitates complex protein-text interactions, paving the way for emerging applications such as text-guided protein design [9, 20] and advanced protein ChatGPT systems [33]. These developments have the potential to significantly advance protein research and engineering.

## Methods

### Protein-function pair construction

For a given protein, we created protein-function pairs using the descriptive information from almost all subsections in the UniProt database [7]. We categorized the subsection information of a protein into two types: sequence-level and residue-level (see Supplementary Table 2). Sequence-level information consists of one or multiple sentences that describe the overall characteristics of the protein, such as its function and catalytic activity. Residue-level information contains phrases describing specific information about certain residues in the protein sequence, which cannot be used directly, such as binding sites and active sites. For residue-level subsections, we employ GPT-4 [39] to generate multiple specific template sentences, thus organizing the information into a coherent sentence. For sequence-level subsections, we also utilize GPT-4 to paraphrase the descriptions and generate multiple alternative sentences to enhance the model’s robustness to textual input. In the end, each subsection in a given protein results in a text sentence, and these sentences are paired with the protein to form protein-function pairs, which are used for training or evaluation purposes.

### Pre-training dataset construction

We first performed a 50% sequence similarity clustering on the human-reviewed Swiss-Prot database [5]. We designated 1000 clusters for validation and another 1000 clusters for testing, using the remaining data as the training set. For each protein, we constructed the protein-function pairs as described above, resulting in a final training set of 14 million protein-function pairs. This high-quality dataset was used to train an initial version of ProTrek. The initial ProTrek model, a 35M version, was pre-trained on 12 NVIDIA 80G A100 GPUs over 100K steps.

Next, we used the initial ProTrek model to score and filter 300 million protein-function pairs from UniRef50 [7]. We retained all protein-function pairs with model scores higher than the average score on Swiss-Prot, resulting in 25 million pairs. These 25 million pairs, combined with the original 14 million pairs, form the final pre-training dataset.

### Pre-training loss function

We adopted InfoNCE loss [43] for protein sequence–structure-function contrastive learning. The InfoNCE loss can be detailed as:

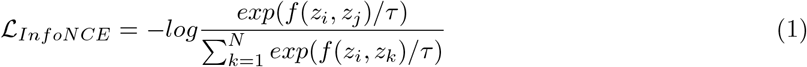

Here, *z*_*i*_ and *z*_*j*_ represent the embeddings of any two modalities. *f* (*z*_*i*_, *z*_*j*_) denotes a similarity score between the embeddings. *N* is the total number of pairs in a batch and *τ* is a learnable temperature parameter. We calculated the InfoNCE loss for the sequence-structure, sequence-function and structure-function pairs from two directions, resulting in 6 contrastive learning functions.

We additionally added 2 Masked Language Modeling (MLM) [40] loss functions to the ProTrek sequence and structure encoder to maintain model’s recognition at the amino acid and 3Di token levels. The training objective is to predict masked tokens by capturing dependencies between masked positions and surrounding context, with the loss function formally described as:

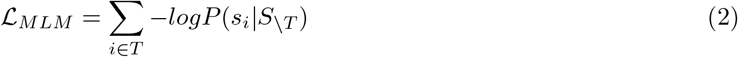

where *T* is a set of token positions to be masked and *S*_*\T*_ represents a protein/structural sequence with tokens at specific positions masked. We formed the final loss function by averaging 2 MLM and 6 InfoNCE loss functions.

### Pre-training setting

We initialized the ProTrek sequence encoder from the ESM-2 650M [19] and the text encoder from the PubMedBERT [15], and we randomly initialized the ProTrek structure encoder as there was no available pre-trained model. For the optimization, we used the DeepSpeed strategy [44] and employed the AdamW optimizer [42], setting *β*_1_ = 0.9, *β*_2_ = 0.98 and we utilized *L*_2_ weight decay of 0.01. We gradually increased the learning rate from 0 to 4e-4 over the first 2000 steps decreased it to 4e-5 using the cosine annealing schedule [41]. The overall training phase lasted approximately 100K steps trained on 20 NVIDIA 80G A100 GPUs. We truncated protein and structural sequences to a maximum of 512 tokens, and truncated the test descriptions to a maximum of 100 tokens. Our total batch size consisted of 1280 protein sequence-structure-function pairs. Additionally, we employed mixed precision training to train ProTrek.

### Protein-function retrieval benchmark

We utilized 4,000 proteins from the Swiss-Prot test set to construct a benchmark for the protein-function task. To evaluate ProTrek’s generalization performance with proteins in other databases, we included 100,000 randomly sampled proteins from UniProt as negative samples in the test set. The textual descriptions of all these proteins were added to construct the text collection. For the protein-to-text search task, the model needed to retrieve the functional descriptions that mostly match the proteins in the Swiss-Prot test set. Conversely, for the text-to-protein task, the model had to find Swiss-Prot proteins that match the query function description (only human-reviewed descriptions were used as query texts) from a set of nearly 104,000 negative sample proteins. We selected the top 33 function subsections based on the number of protein-text pairs for evaluation. This approach aims to mitigate the bias impact of subsections with fewer pairs.

### Detecting proteins with similar functions

For ProTrek and all baseline methods, we conducted an all-versus-all search using the same Swiss-Prot test dataset consisting of 4,000 proteins and compared their performance for finding proteins of the same GO annotation. for a given query protein, *G*_*q*_ represents the set of GO annotations associated with that protein. Similarly, for each hit retrieved from the database, *G*_*h*_ represents the set of GO annotations assigned to that hit. We defined a hit as correct if there was at least one common GO annotation between *G*_*q*_ and *G*_*h*_, denoted as *G*_*q* ∩_ *G*_*h*_ ≠ ∅. To generate the y-axis values in Fig. 2c, we counted the number of correct hits across all query proteins in the test set and summed them up.

### Mean average precision

Mean Average Precision (MAP), a commonly used metric for information retrieval tasks, is the Average Precisions (APs) for each query. For a given query, AP is calculated by the following formula:

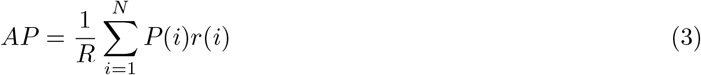

where *R* is the number of relevant results and *N* denotes the total number of results. For the *i*^*th*^ result, *P* (*i*) is the precision at the *i*-th position in the ranked list, and *r*(*i*) indicates whether the result is relevant to the query (1 if relevant and 0 if not relevant). The MAP is calculated by averaging the APs from all queries:

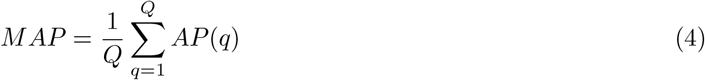

### Deep learning baseline models

For protein-function searches, we compared ProTrek to ProtST-ESM-1b from ProtST [35] and Prot-BERT BFD-512-1e-5-1e-1-text-512-1e-5-1e-1-InfoNCE-0.1-batch-9-gpu-8-epoch-5 from ProteinDT [20]. For the downstream protein tasks, we used the ESM-2 650M version [19] and ProtST-ESM-1b as baselines.

### Downstream task fine-tuning

We strictly followed SaprotHub’s fine-tuning settings [27], including dataset pre-processing, splitting, finetuning hyper-parameters, etc. We fine-tuned all model parameters until convergence and selected the best checkpoints based on their performance on the validation set.

### Tool execution commands

We ran each alignment-based tool according to its standard usage example:

### Foldseek

We used Foldseek with the **ef4e960ab84fc502665eb7b84573dfff9c2aa89d** version. The command line was executed with default parameters: **foldseek easy-search pdb dir targetDB aln.m8 tmpFolder**

### MMseqs2

We used MMseqs2 with the **edb8223d1ea07385ffe63d4f103af0eb12b2058e** version. The command line was executed with default parameters: **mmseqs easy-search seqs.fasta targetDB alnRes.m8 tmp**

### BLASTP

We used BLASTP with the **Protein-Protein BLAST 2.15.0+** version. We ran BLASTP from the command line: **blastp -query seqs.fasta -db db -outfmt 6 -out blastp result**

### DIAMOND

We used DIAMOND with the **v2.1.9.163** version. We ran DIAMOND in very-sensitive mode following the standard case: **diamond blastp -q seqs.fasta -d db -o result.tsv –very-sensitive -k 0**

## Data availability

The pre-trained ProTrek weights can be downloaded from https://huggingface.co/westlake-repl/ProTrek_650M_UniRef50. All pre-computed protein sequence, structure and text embeddings for Swiss-Prot database are available at https://huggingface.co/datasets/westlake-repl/faiss_index_ProTrek_650M_UniRef50 (will release protein embeddings for other database soon). The structural 3Di sequences are available at https://github.com/steineggerlab/foldseek. The text descriptions are available at https://www.uniprot.org/

## Code availability

ProTrek is open-sourced under the MIT license. The code repository is available at https://github.com/westlake-repl/ProTrek. The ProTrek webserver is located at https://huggingface.co/spaces/westlake-repl/Demo_ProTrek_650M_UniRef50. The ColabProTrek is available at https://colab.research.google.com/drive/1On2xQU0d7351bIBgZpz2T0VUp2gZium0?usp=sharing.

## Acknowledgements

We thank Sergey Ovchinnikov and Chentong Wang for their valuable suggestions to improve this paper.

## Supplementary information

**Supplementary Table 1.**
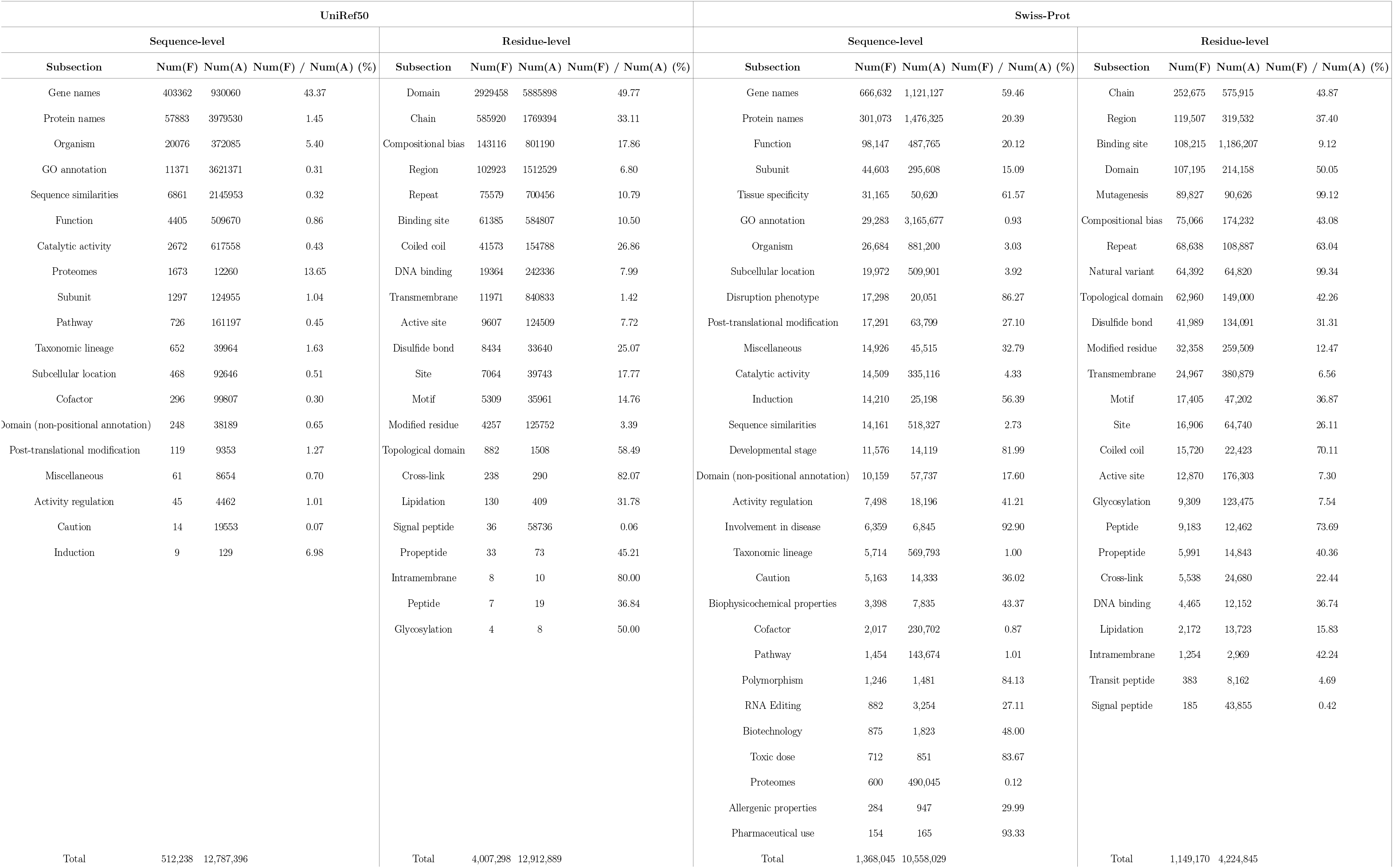
Statistics of the pre-training dataset. **Num(F)** represents the number of unique descriptions, while **Num(A)** indicates the total number of descriptions, including replicates.

**Supplementary Table 2.**
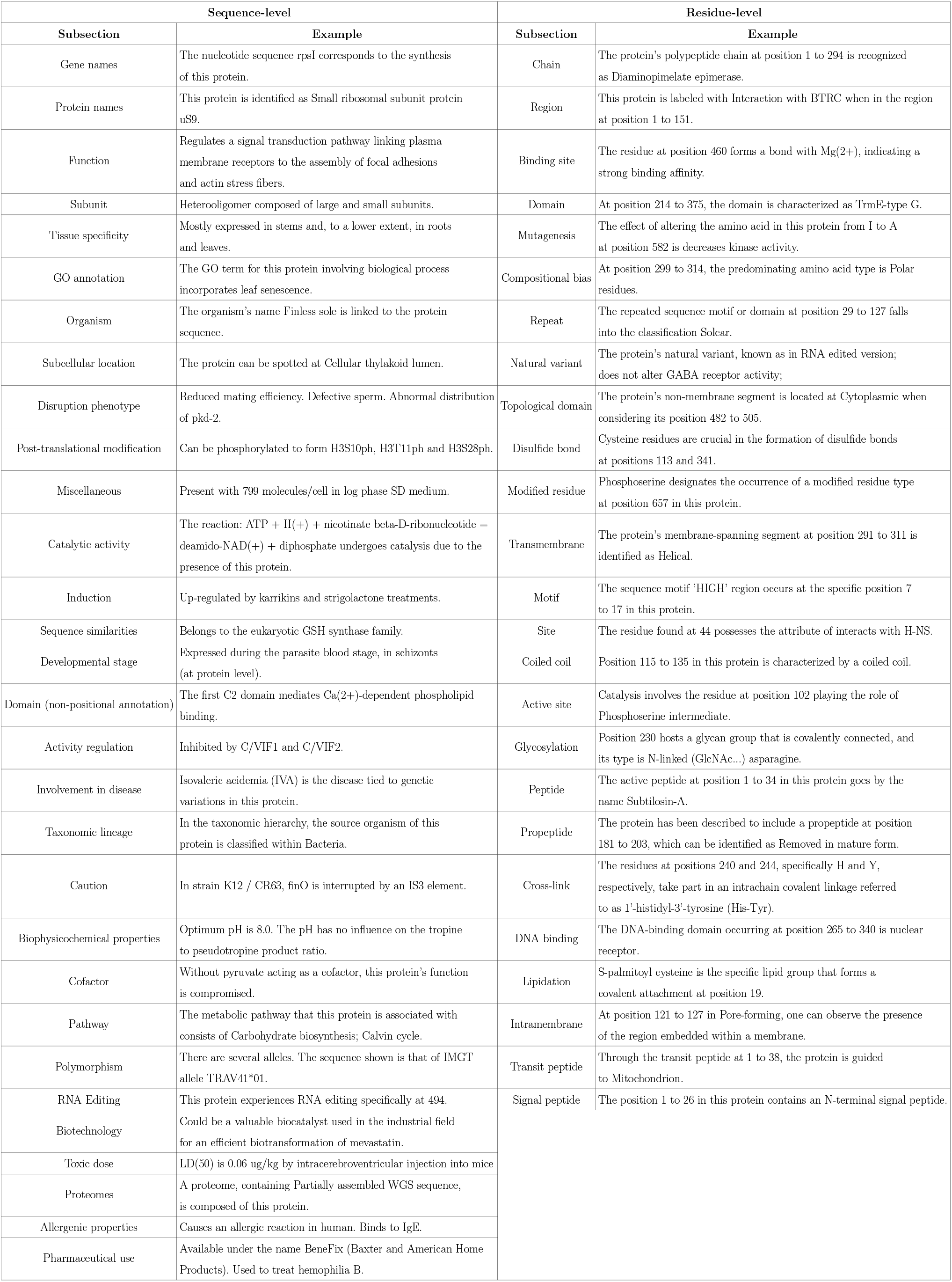
Examples of functional descriptions for each subsection. The examples in the residue-level subsections are created by filling in the keywords (e.g. position) in the templates.

**Supplementary Fig. 1.**
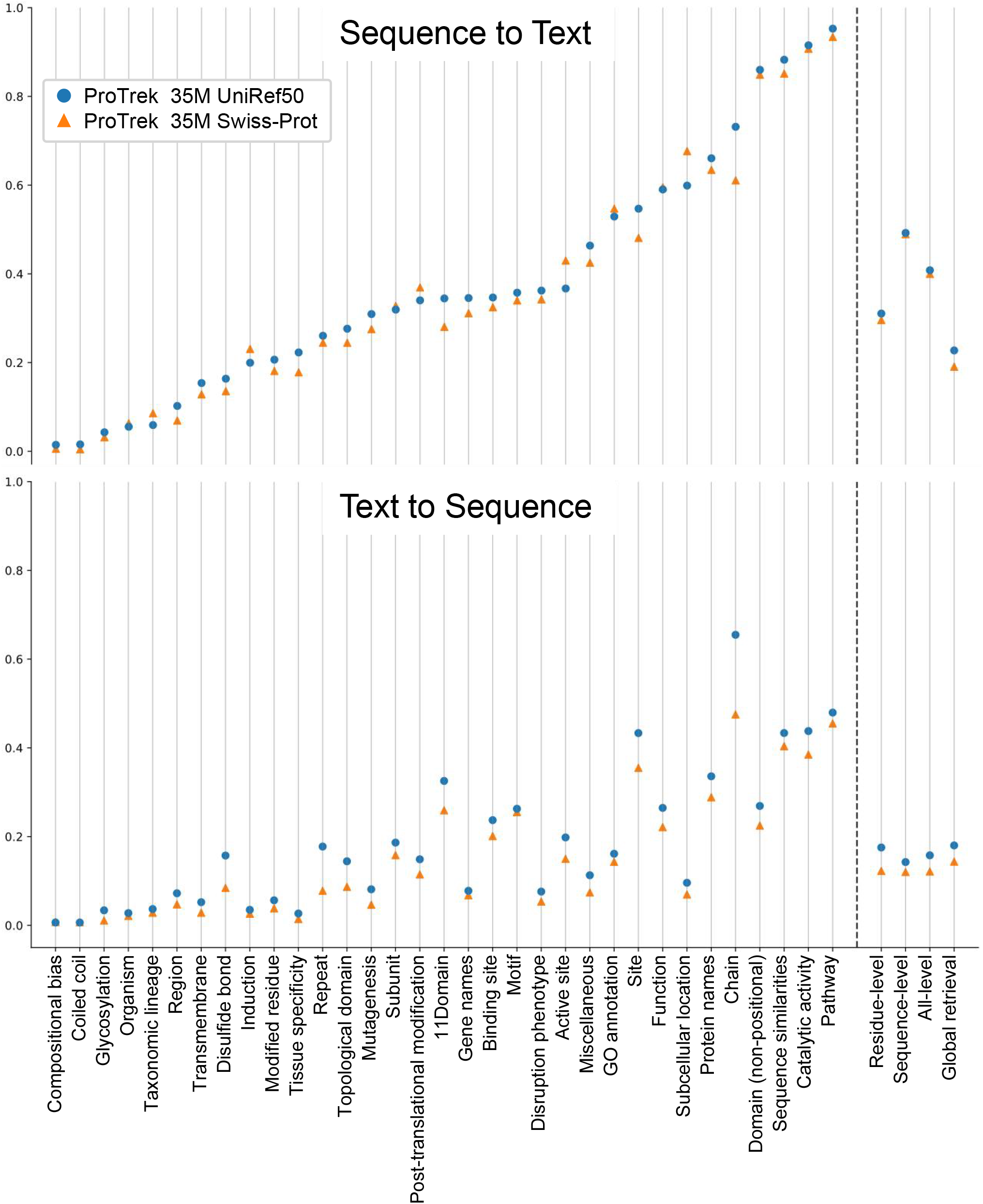
Comparison of protein-text retrieval between models trained on different datasets. “ProTrek 35M Swiss-Prot” was trained on Swiss-Prot data only and “ProTrek 35M UniRef50” was trained on a mixture of Swiss-Prot and UniRef50 data.

**Supplementary Fig. 2.**
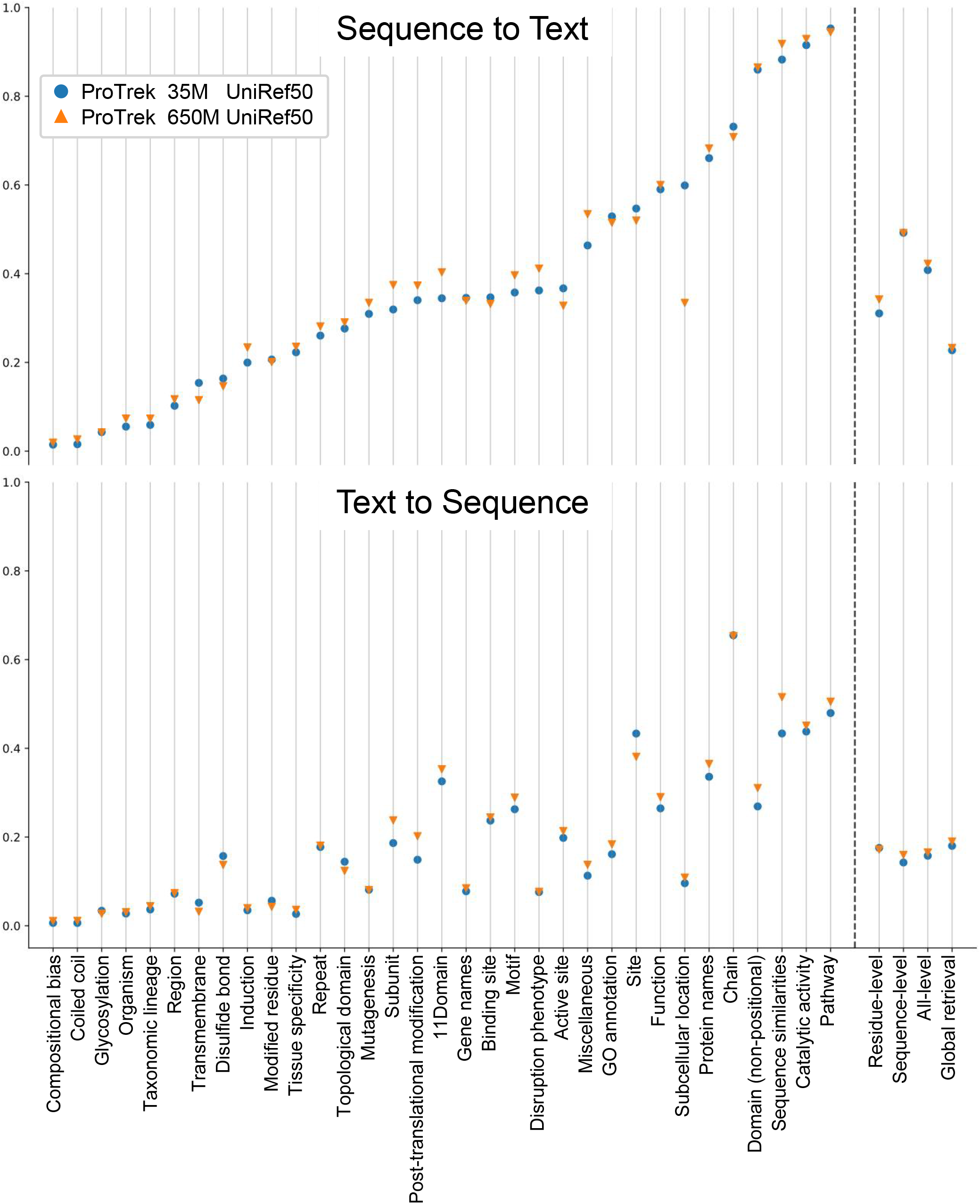
Performance comparison of the ProTrek 35M and ProTrek 650M.

**Supplementary Fig. 3.**
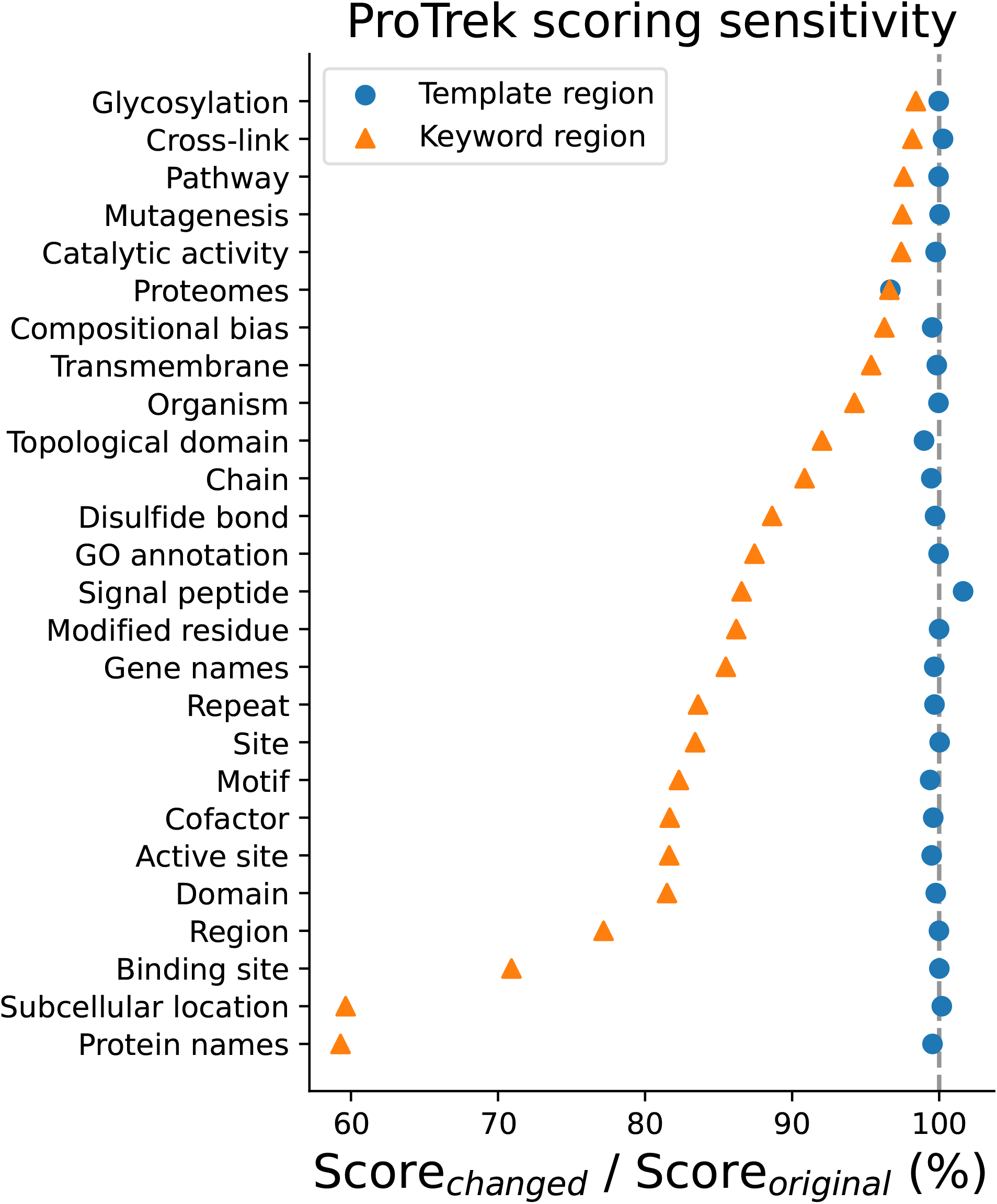
Scoring sensitivity for keyword regions and template regions in residue-level functional descriptions. Initially, we computed the similarity score “Score_*original*_” between a protein sequence and a textual description. Subsequently, we removed each word in the description and calculated a new similarity score with the protein. For the template region (highlighted with a blue circle), we averaged all the new scores whose deleted words were in the template region, denoted as “Score_*changed*_”. Similarly, the “Score_*changed*_” of the keyword region (indicated by an orange triangle) was calculated by averaging all the new scores whose deleted words were in that region.

**Supplementary Fig. 4.**
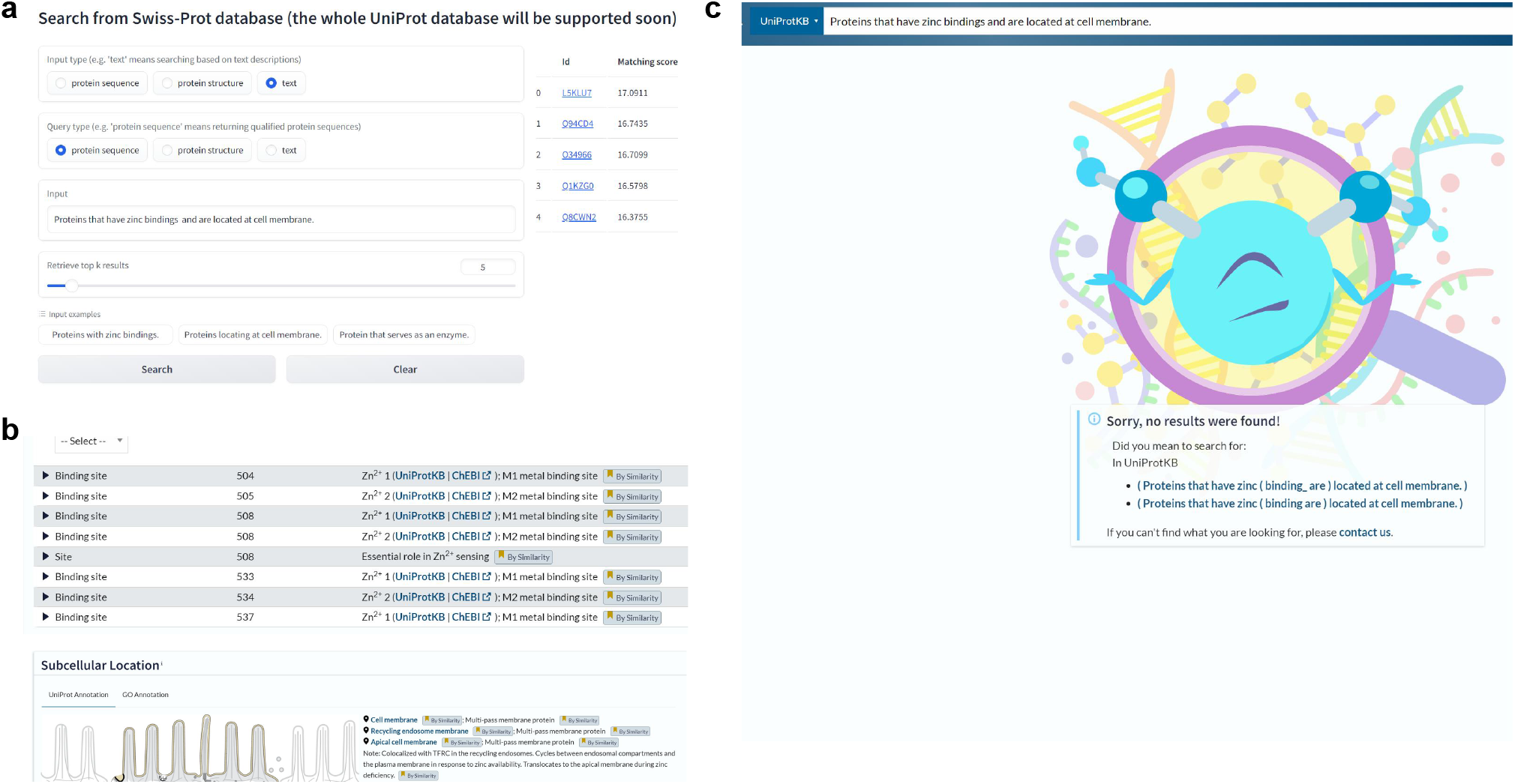
A case study using ProTrek to identify proteins with a long functional description. **a**, Using ProTrek demo to search proteins with the long functional description “Proteins that have zinc bindings and are located at cell membrane.” **b**, “L5KLU7,” a human-reviewed protein, is returned as the top-ranked result by ProTrek, matching the corresponding description exactly. **c**, Using the same description to search against the UniProt database. The UniProt database only supports keyword searches and did not return any results for the long natural sentence.

## Footnotes

1 Initial startup may take a few minutes due to sleep mode. The ProTrek server will support larger databases like BFD (https://bfd.mmseqs.com/), MGnify (https://www.ebi.ac.uk/metagenomics), and OMG [8], containing billions of protein sequences.

